# Coessentiality and cofunctionality: a network approach to learning genetic vulnerabilities from cancer cell line fitness screens

**DOI:** 10.1101/134346

**Authors:** Traver Hart, Clara Koh, Jason Moffat

## Abstract

Genetic interaction networks are a powerful approach for functional genomics, and the synthetic lethal interactions that comprise these networks offer a compelling strategy for identifying candidate cancer targets. As the number of published shRNA and CRISPR perturbation screens in cancer cell lines expands, there is an opportunity for integrative analysis that goes further than pairwise synthetic lethality and discovers genetic vulnerabilities of related sets of cell lines. We re-analyze over 100 high-quality, genome-scale shRNA screens in human cancer cell lines and derive a quantitative fitness score for each gene that accurately reflects genotype-specific gene essentiality. We identify pairs of genes with correlated essentiality profiles and merge them into a cancer coessentiality network, where shared patterns of genetic vulnerability in cell lines give rise to clusters of functionally related genes in the network. Network clustering discriminates among all three defined subtypes of breast cancer cell lines (basal, luminal, and Her2-amplified), and further identifies novel subsets of Her2+ and ovarian cancer cells. We demonstrate the utility of the network as a platform for both hypothesis-driven and data-driven discovery of context-specific essential genes and their associated biomarkers.

## Introduction

The concept of synthetic lethality, where one gene becomes essential in the presence of a second gene’s mutation or loss of function, has long been recognized as a powerful strategy to finding candidate therapeutic targets for cancer [1]. Recently, several promising leads for chemotherapeutic targeting were discovered by identifying likely synthetic lethal gene pairs where one member of the pair is frequently co-deleted alongside a neighboring tumor suppressor gene – an approach called collateral lethality [2]. In glioblastoma, for example, the glycolytic gene enolase 1 (ENO1) is frequently deleted, rendering those cells specifically dependent on the gene’s homologue, ENO2 [2]. Similarly, in pancreatic cancer, malic enzyme 2 (ME2), which converts malate to pyruvate in the mitochondria, imparts a selective dependency on its paralog ME3 when ME2 is co-deleted with tumor suppressor SMAD4 [3].

The best known clinical application of synthetic lethality is the emergent sensitivity to PARP inhibitors discovered in BRCA1 and BRCA2 mutant cells [4-6]. The FDA has recently expanded the clinical application of the PARP inhibitor olaparib beyond BRCA1/2-mutant ovarian cancer to patients with BRCA1/2 or ATM-mutated advanced prostate cancer. The addition of ATM-mutant backgrounds is the result of an important trend in preclinical research regarding olaparib inhibitor efficacy: tumors (and cell lines) deficient in any of several components of the homologous recombination (HR) mediated DNA double strand break repair machinery are highly dependent on alternative repair pathways mediated by PARP [7, 8]. Mutations in more than a dozen genes involved in HR and other DNA damage response pathways are also associated with reliance on PARP and, in turn, increased sensitivity to olaparib [9, 10].

Several genetic screening approaches are currently underway to systematically define the network of synthetic lethal relationships in human cells, in order to refine our knowledge of functional genomics and to exploit specific interactions for cancer targeting. Digenic knockout screens in human cells using dual-gRNA CRISPR/Cas9 constructs have demonstrated the potential of targeted pairwise gene knockout screens [11, 12], but scalability is a major issue and technology advances will be required before this strategy can be employed on a genomic scale.

A second approach, whole-genome screens across a panel of isogenic “query gene” knockout cell lines, offers considerable advantages for functional genomics: a single, genome-scale perturbation library can be developed and re-used across a large number of experiments, allowing relatively easy data integration. This approach has been used with great success in yeast: a massive survey of 23 million double mutants defined a global map of genetic interactions [13, 14] and demonstrated that genes with correlated genetic interaction profiles across a panel of query strains were often involved in the same biological processes, enabling functional characterization of previously uncharacterized genes. Such a strategy holds great promise for human functional genomics studies but may be less applicable for cancer targeting, as tumor-relevant synthetic lethals are often context dependent and not generalizable from isogenic screens [10, 12, 15]. Indeed the dynamics of genetic interaction networks are also an area of active research [16, 17].

The third strategy involves genome-scale perturbation screens across a large panel of genetically diverse cancer cell lines. The integrated analysis of such data would reveal genes that are consistently essential across similar cell lines (e.g. those sharing a common driver oncogene), helping to address the generalizability issue in isogenic screens [18]. It would aid the identification of genetic vulnerabilities that are specific to a given genetic background or subtype by demonstrating their nonessentiality outside of that subtype. Also, such a study would carry its own internal controls, as functionally related genes should show correlated patterns of essentiality across sufficiently diverse backgrounds.

In this study, we describe the integrated analysis of a large compendium of genetic perturbation screens. Using over 100 genome-scale, pooled-library shRNA screens from breast [19], ovarian [20], and pancreatic [21] cancer cell lines, all conducted with a common shRNA library and using similar experimental designs, we show that optimizing for co-functionality reveals a gene coessentiality network whose structure is driven by the shared genetic vulnerabilities of the cell lines. We demonstrate that clusters of coessential genes define known as well as novel subgroups of cell lines, and show how the network can be integrated with tumor molecular data to predict known and novel drug targets and their biomarkers. This approach demonstrates how the integrated analysis of noisy cell line screens can be used for hypothesis-driven as well as data-driven discovery.

## Results

### Generating the coessentiality network

A fundamental insight from the systematic survey of genetic interactions in yeast is that if two genes have similar interaction profiles across the same panel of genetic backgrounds, they are likely to be involved in the same biological process [22]. We reasoned that an analogous principle should hold in human cell lines: that a gene’s perturbation-induced fitness defect likely varies across different genetic backgrounds, yielding a gene “essentiality profile,” and that genes with correlated essentiality profiles (“coessential genes”) should be involved in the same cellular functions. We hypothesized that, given a high-quality set of perturbation fitness screens across a sufficiently diverse set of tumor genetic backgrounds and tissues of origin, patterns of covariation in essentiality profiles that correspond to known tissues or subtypes might reveal new context-specific essential genes that could potentially serve as novel therapeutic targets. Moreover, we reasoned that the cell lines driving these patterns of covariation might reveal novel genetic or phenotypic subtypes based on shared genetic vulnerabilities.

To this end, we re-analyzed a set of genome-scale pooled library shRNA screens in human cell lines [19-21]. All the screens were conducted with an shRNA library containing 78,432 lentiviral-encoded shRNA hairpins targeting 16,056 human Refseq protein-coding genes, facilitating comparisons across screens. We applied the BAGEL algorithm to generate a log Bayes Factor of gene essentiality for each gene in each cell line, resulting in a matrix of scores for 12,913 genes across 112 cell lines (Figure 1a), with matching RNAseq gene expression profiles (Supplementary Tables 1-3).

**Figure 1.**
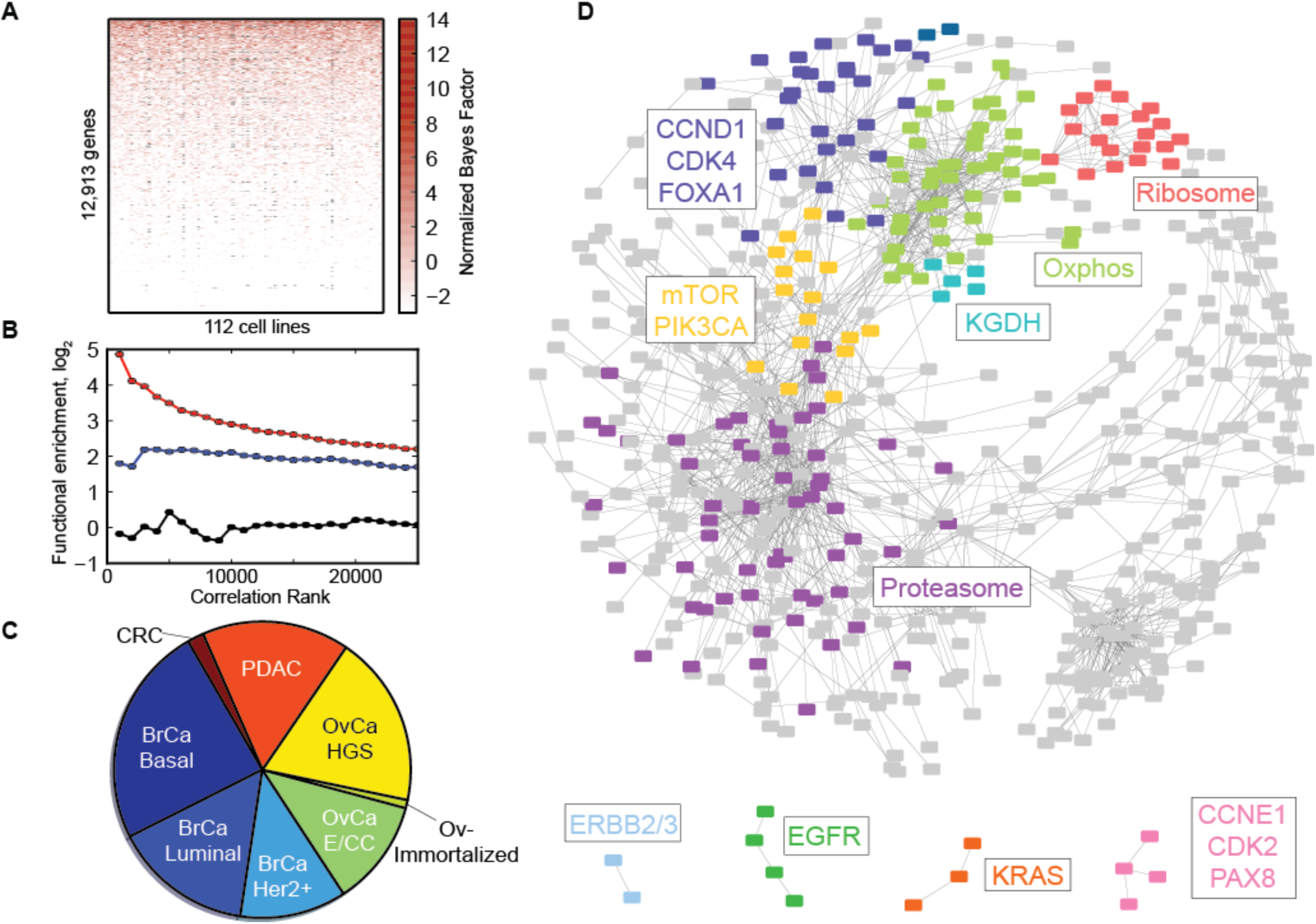
The coessentiality network. (A) Raw data is a matrix of Bayes Factors for >15,000 genes across >100 cell line screens. (B) Correlation of essentiality scores across screens vs. cofunctionality (y-axis). Black, raw data gives no signal. Blue, normalized BFs improve cofunctionality signal. Red, filtering for high-confidence essential genes yields 30-fold enrichment for co-functionality. (C) Final data set consists of 112 screens from breast, ovarian, pancreatic, and colorectal cancer cell lines. (D) The cancer coessentiality network of 2,883 genes, with major clusters highlighted and annotated with characteristic genes/systems.

We measured the functional coherence of this dataset by calculating the Pearson correlation coefficient of the fitness profiles of all pairs of genes, rank-ordering gene pairs by correlation coefficient, binning into groups of 1,000 pairs, and measuring the cumulative log likelihood of shared KEGG terms in each bin ([23]; see Methods). Overall, the raw Bayes Factor matrix yielded no relationship between fitness profile correlation and functional interaction (Figure 1b, black).

After observing major technical (i.e. non-biological) sources of variation in the data, we applied a variety of filtering and normalization measures in an attempt to maximize the functional enrichment of highly correlated genes. Initially, we normalized each gene’s BF score by the number of hairpins used to generate the score (see Methods) and quantile normalized the matrix. This yielded a fourfold enrichment in co-functionality across highly correlated pairs (Figure 1b, blue). Then, based on observations that shRNA screens are most accurate when targeting high-expression genes [24], we included only genes which were called essential (normalized BF>2) in at least one cell line where the gene also showed mRNA expression above median for that sample (see Methods). This yielded a final data set of 2,883 genes in 112 cell lines (Figure 1c and Supplementary Table 4). Genes with highly correlated essentiality profiles in this filtered data set showed nearly 30-fold enrichment for involvement in the same biological process, validating our hypothesis that human essentiality profiles are analogous to yeast genetic interaction profiles (Figure 1b, red). Hereafter, we refer to each gene’s hairpin- and quantile-normalized Bayes Factor in this matrix as its *essentiality score*, where an essentiality score > 2 is considered a high-confidence hit.

We then merged the gene pairs with the highest correlations (adjusted P-value < 0.001, corresponding to a Pearson correlation coefficient of 0.457 across 112 samples) into a network where nodes represent genes and edges connect genes with highly correlated essentiality profiles. Most of the gene pairs merged into one giant connected component, with 1,484 edges connecting 564 genes (Figure 1d and Supplementary Table 5). We clustered the network using mcl [25], an implementation of Markov clustering, and observed that many clusters showed high functional coherence, as would be expected from a network trained on maximizing the presence of co-functional gene pairs. In particular, the ribosome and proteasome appeared in distinct clusters in the coessentiality network (Figure 1d, red and purple clusters; Supplementary Table 6).

### Clusters in the coessentiality network are defined by distinct cell types

Though the presence of functionally coherent clusters serves as a useful positive control for our approach, our motivating hypothesis was that other clusters of coessential genes might reveal subtype-specific patterns of genetic vulnerability. Indeed, one of the largest clusters in the network contained well-characterized oncogenes CCND1, CDK4, and FOXA1, known to be specific to breast cancers of luminal subtype (Figure 1d, dark blue). We extracted the 29 genes in this cluster from the essentiality matrix and performed hierarchical clustering on the cell lines. As expected, luminal and Her2-amplified breast cancer cell lines clustered together with higher essentiality scores across this panel of genes (Figure 2a). A smaller cluster containing only ERBB2 and ERBB3 (Figure 1d) segregates Her2-amplified cell lines (Figure 2b). Interestingly, a third cluster further subdivides the Her2-amplified cell lines. The five genes in the cluster include all three subunits of the mitochondrial alpha-ketoglutarate dehydrogenase complex (DLD, DLST, OGDH; Figure 2c), a key component of the TCA cycle. This complex is strongly essential in five Her2+ cell lines but nonessential in 8 others, suggesting the presence of further metabolic subtypes within the recognized Her2+ subtype [26]. A recent report indicates that dependency on αKGDH is driven by PIK3CA mutations [27]. However, the differential sensitivity to perturbation in our network is not correlated with PIK3CA mutation status: 3/5 sensitive cell lines and 4/8 insensitive cell lines carry oncogenic PIK3CA mutations (P=0.59, Fisher’s exact test). The emergence of known, well-defined subtypes from the gene clusters in the coessentiality network validates the effectiveness of this approach, and the indication of even more fine-grained subtyping indicates the power of the coessentiality network to discover novel relationships between cell lines.

**Figure 2.**
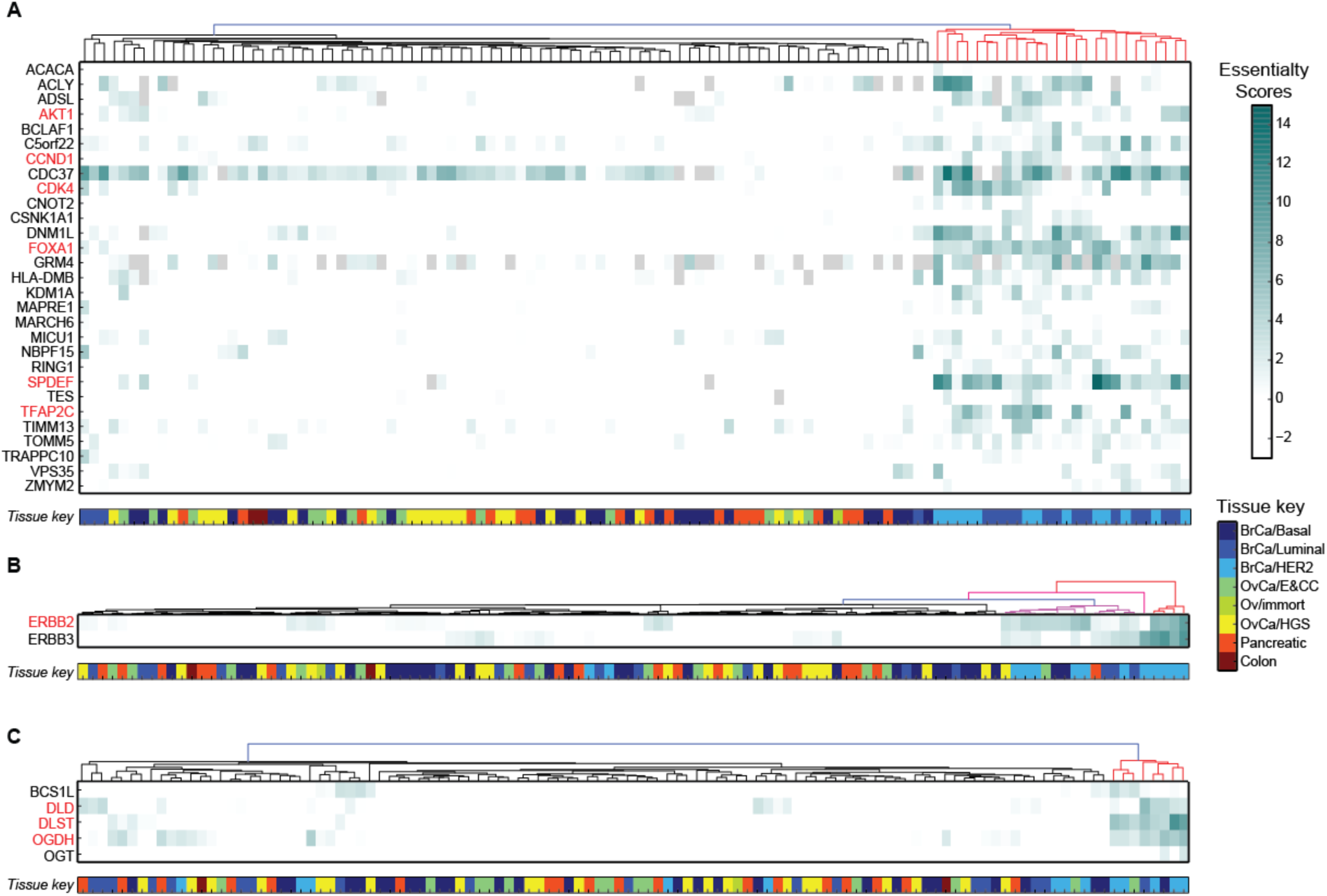
Cancer subtypes drive coessential clusters. (A) Large cluster containing known oncogenes (red; e.g. CKD4, CCND1, FOXA1) is driven by subtype specific essentiality in BrCa/Luminal and BrCa/Her2+ cell lines (tissue key, bottom). (B) Smaller cluster containing ERBB2 and ERBB3 differentiates BrCa/Her2+ cell lines. (C) Cluster with all three subunits of alpha-ketoglutarate dehydrogenase (KGDH) complex divides BrCa/Her2+ cell lines into two subgroups.

Among the largest clusters in the network is a group of 50 genes, of which 45 encode proteins which operate in the mitochondria (P<5 × 10^−44^, 11-fold enrichment; Figure 1e, green cluster). These genes are highly enriched for genes encoding subunits of the electron transport chain complexes, as well as other proteins involved in processes required for ETC complex biogenesis (e.g. the mitochondrial ribosome). Hierarchical clustering the cell lines according to their essentiality profiles across these genes yields three dominant clusters, two of which are clearly associated with higher essentiality scores in this set of genes (Figure 3a). These two sets of cell lines collectively contain 11 of 13 Her2-amplified breast cancer cell lines as well as 12 of 35 ovarian cancer cell lines (9 of 21 HGSOC and 3 of 14 E/CC cell lines). To validate this cluster, we selected five ovarian cancer cell lines predicted to be oxphos-sensitive and nine predicted to be oxphos-insensitive and treated them with the Complex I inhibitor rotenone. All five oxphos-sensitive cell lines showed lower viability in the presence of rotenone than the nine insensitive lines (Figure 3b).

**Figure 3.**
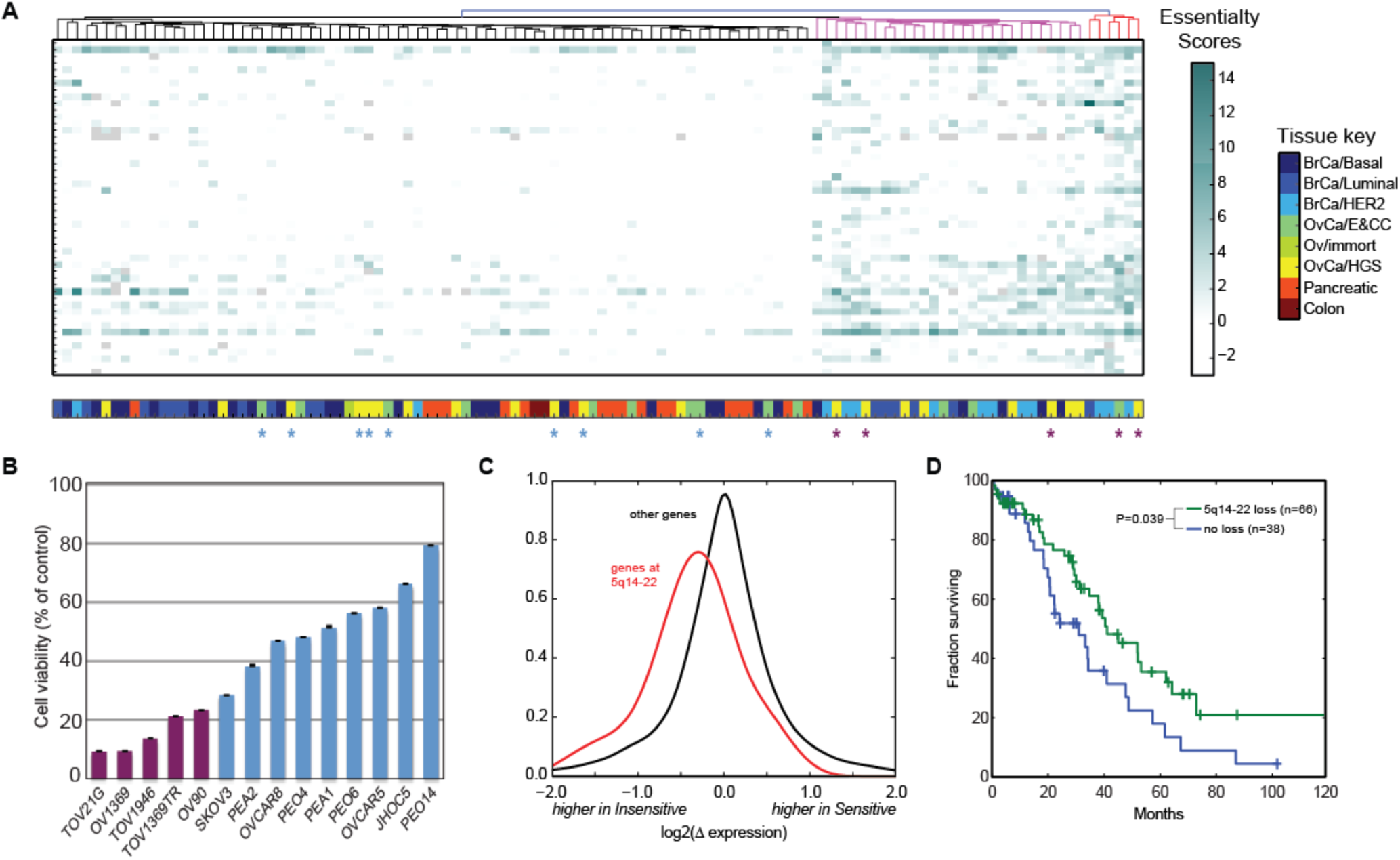
Oxphos cluster. (A) A large cluster of mitochondrial genes is preferentially essential in BrCa/Her2+ and a subset of OvCa cell lines. A subset of OvCa cell lines (blue, purple stars) were selected for further validation. (B) Relative viability of OvCa lines in the presence of Rotenone. (C) Differential expression of OvCa/HGS oxphos sensitive and insensitive cells showed strong bias at 5q14-22 locus, consistent with copy number aberration. (D) TCGA data shows 5q14-22 copy loss provides survival benefit in HGSOC patients.

Complex I is gaining attention as a candidate therapeutic target in cancer, in particular as a target of the widely prescribed antidiabetic drug metformin[28, 29]. Identifying a biomarker of Complex I sensitivity could have major clinical relevance. To explore whether our observed difference in phenotype among similar cancer cell lines might reflect some identifiable molecular difference in tumors, we searched for functional phenotypes or biomarkers that might segregate the cell lines. Surprisingly, we found no significant functional enrichment among differentially expressed genes (Supplementary Table 6), including no difference among gene sets related to oxidative phosphorylation or mitochondrial function. A subsequent analysis based on genomic coordinates of differentially expressed genes did, however, reveal several loci that are candidates for copy number aberrations (Supplementary Table 7). In particular, genes located at 5q14-22 show significantly lower expression in the oxphos-sensitive cells, suggesting a genomic copy loss at that locus (Figure 3c). In TCGA data from ovarian tumors [30, 31], tumors with a heterozygous or homozygous copy loss at this locus show a significant increase in median survival, especially for tumors of the mesenchymal subtype (41 vs. 31 months; P=0.039, log-rank test; Figure 3d). The survival difference appears to be limited to the mesenchymal subtype: including all ovarian tumors (n=481) improves the P-value (P=0.0049) but reduces the median survival difference to 2.5 months, while no difference is indicated for proliferative (P=0.12), immunoreactive (P=0.82), or fallopian subtypes (P=0.85). This example highlights an indirect approach to discovering tumor subtypes of potential clinical relevance: identifying phenotypic differences between groups of cell lines, finding biomarkers that segregate those phenotypes, and applying those biomarkers to primary tumor data.

### Core essentials are more sensitive to perturbation at lower expression levels

The matrix of gene essentiality scores holds utility beyond the correlation network described. We examined the relationship between gene expression and essentiality by calculating the Pearson correlation coefficient between a gene’s essentiality profile and its expression profile across the same samples. Across the 2,883 essential genes in the final data set, the distribution of correlation coefficients is roughly normally distributed and centered at zero (Figure 4a), indicating no general relationship between variation in expression and variation in essentiality. However, we observed two notable exceptions to this general trend. First, “core essential” genes—genes expected to be essential across all cell lines [32]—show a strong bias toward negative correlation, indicating that lower expression implies a greater sensitivity to perturbation (Figure 4a, orange curve). This is broadly consistent with studies of variation in gene expression across *C. elegans* strains [33] as well as the increasing body of work suggesting heterozygous copy loss of core essential genes in cancer cells—and the commensurate lower gene expression—increases sensitivity to drugs targeting those genes and pathways [2, 34-36]. Second, at the other end of the spectrum, genes with highly positive expression/essentiality correlation tend to be tissue-specific essential genes. Of the nine genes with correlation coefficient >= 0.4 (Figure 4a, inset), all showed a strong tendency toward essentiality in a specific subtype, including previously mentioned SPDEF and FOXA1 in luminal-subtype breast cancer cells, ERBB2 and estrogen receptor gene ESR1 in Her2+ and luminal-subtype cells, respectively, and FUBP1 and PAX8 in ovarian cancer cells.

**Figure 4.**
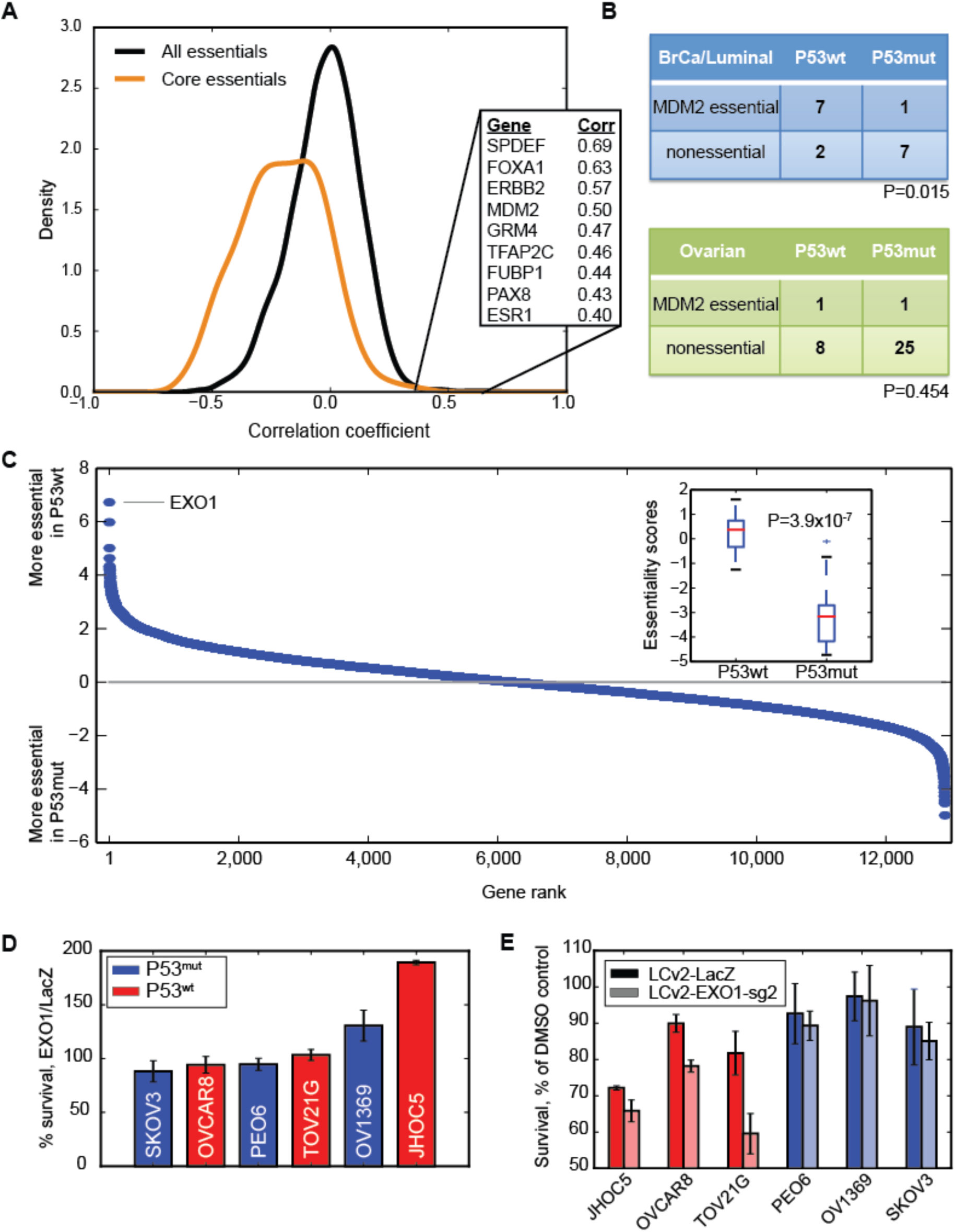
(A) Most essential genes show no correlation between expression and essentiality (black). Core essentials show negative correlation (orange). The small number of positively correlated genes are highly enriched for subtype-specific essential genes (inset). (B) MDM2 is essential in P53wt cell lines in BrCa (top) but not in OvCa (bottom). (C) Exonuclease EXO1 is the top differentially essential gene in OvCa lines expressing WT vs mutated P53, but is not a highly essential gene. (D) EXO1 knockout does not kill P53 mutant (red) or P53 wildtype (blue) ovarian cancer cells. (E) Survival in the presence of nutlin 3a. EXO1 knockout sensitizes P53 wildtype cells but not P53 mutant cells to MDM2 inhibition.

### EXO1 depletion sensitizes OV/ECC cells to MDM2 inhibition

We expected that the expression-dependent essentiality of MDM2, a gene whose major role in cancer is the suppression of *P53* protein expression, would be associated with subtypes that are not characterized by genetic suppression of pro-apoptotic TP53 activity, e.g. by mutation, deletion, or methylation. We found this to be true for BrCa/luminal cell lines (P=0.015; Fisher’s exact test) but, surprisingly, not true for ovarian cancer cell lines, where endometrioid and clear cell subtypes are often characterized by wildtype TP53 (P=0.454; Figure 4b).

To explore the possibility that these cells rely on different DNA damage response pathways, we compared the essentiality scores in ovarian cancer cell lines with (n=21) and without (n=7) TP53 mutations, after excluding 7 cases with low TP53 mRNA expression (Figure 4C). The top hit was EXO1, an exonuclease involved in DNA mismatch repair and double strand break repair [37, 38]. While P53wt and P53mut ovarian cancer cell lines showed a marked difference in sensitivity to perturbation of EXO1 by shRNA (P=3.9 × 10^−7^, t-test; Figure 4c, inset), it is worth noting that even the EXO1-sensitive lines did not meet our threshold of high-confidence hits (essentiality score >=2). This is consistent with the fact that CRISPR-mediated EXO1 knockout in P53wt cells did not elicit a severe fitness defect (Figure 4d). However, EXO1 deletion did restore sensitivity to MDM2 inhibition: EXO1^null^ P53wt cells are more sensitive to MDM2 inhibitor nutlin-3a than their EXO1^wt^ parental strains (Figure 4e), supporting the existence of a synthetic interaction between *p53* mutation state and EXO1 in this cell type and further confirming the utility of the coessentiality network to identify novel context-specific genetic interactions.

## Discussion

A key insight from the systematic survey of yeast genetic interactions is that genes which operate in the same biological processes tend to have similar profiles of genetic interactions across a diverse panel of query strains; that is, they show the same patterns of fitness defects across different genetic backgrounds. We applied this concept to the analysis of genetic perturbation screens in human cell lines, using the BAGEL-derived Bayes Factor as a fitness score. Initially, the approach did not appear to work, as correlated gene fitness profiles showed no enrichment for co-functionality. However, after filtering the data to include only high-quality screens and high-confidence essential genes, the picture came into focus, with highly correlated genes showing nearly thirty-fold enrichment for shared biological process annotations. This preliminary result further illustrates the value of the BAGEL algorithm in offering a semi-quantitative measure of gene knockdown fitness as well as the utility of the approach outlined in [32] to distinguish high quality screens from those that should be removed from downstream analyses.

We combined the highly correlated gene pairs into a network of genes, where genes are connected by an edge if they show a correlated fitness profile across the panel of 112 cell lines. Clustering this network revealed groups of genes, some of which operate in well-annotated biological pathways—e.g. the ribosome and proteasome clusters—while other genes were grouped together based on patterns of covariation in fitness across the different cellular contexts included in the network. Clusters identifying genes specifically essential in luminal and Her2-amplified breast cancer cell lines, for example, were readily identified in the network, validating our approach. Furthermore, among the novel clusters was a large cluster of genes related to mitochondrial function that segregate ovarian cancer cell lines into oxphos-sensitive and oxphos-resistant classes. This latter group shows a molecular signature represented by chromosomal copy loss in the 5q14-22 region that, in turn, offers a survival advantage for ovarian cancer patients. Thus, the coessentiality network offers a method of identifying both novel differentially essential genes across known subtypes but also a way to discover new subtypes, with possible clinical relevance, from a large body of shRNA knockdown data.

The matrix of essentiality scores we generated offers utility beyond the network approaches described. We measured the correlation between these normalized fitness scores and mRNA expression levels of the same gene, which revealed three distinct subgroups. Tissues-specific essentials show high expression/fitness correlation but are rare among the subtypes that were assayed here. Core essentials, in contrast, showed an increasing sensitivity to perturbation at lower expression levels. This is generally consistent with the concept of genomic copy loss of essential genes leading to a possible therapeutic window for drug targeting, and extends the pool of candidate targets to include all core essential genes (i.e. those essential in virtually all backgrounds). Finally, the bulk of the signal is that there is no signal: in general essential genes showed no correlation between their expression level and their knockdown fitness. This has potentially major implications for efforts to analyze the large corpus of tumor molecular data generated by, for example, The Cancer Genome Atlas. If the majority of context-dependent essential genes show no relationship between fitness and expression level, or other molecular signatures, then this large class of potential therapeutic targets may be invisible to analyses of tumor molecular data alone.

Fortunately, large-scale screening of cancer cell lines using CRISPR/Cas9 gene knockout libraries is underway, providing a systematic view of genetic vulnerabilities in cancer cell lines [24, 39-42]. We and others have shown that CRISPR/Cas9 fitness screens can provide an enormous improvement in both sensitivity and specificity over shRNA screens. The application of the coessentiality network approach described here to a large body of diverse CRISPR screens should reveal even deeper insight about context-specific vulnerabilities and their biomarkers, as well as the functional genomics of cancer more generally.

## Acknowledgments

We wish to thank members of the Moffat and Hart labs for helpful discussions. TH was supported by MD Anderson Cancer Center Support Grant P30 CA016672 (the Bioinformatics Shared Resource) and the Cancer Prevention Research Institute of Texas (CPRIT) grant RR160032. This work was supported from grants funded by the Canadian Institutes for Health Research to JM (MOP-142375) and the Ontario Research Fund to JM (ORF-RE7-037). JM is a Canada Research Chair (Tier 2) in Functional Genetics.

## Methods

### Primary data processing

Data from shRNA screens in pancreatic, ovarian, and breast cancer cell lines was downloaded from the Donnelly-Princess Margaret Screening Centre (formerly COLT [43]) at dpsc.ccbr.utoronto.ca. The shRNA and RNA-seq data are from three published studies of shRNA screens [19-21].

Each screen consists of a reference timepoint (T0) and one to two experimental timepoints (T1, T2), typically assayed in triplicate using custom microarrays. See [21] for experimental details. shRNA hairpins (hereafter ‘hairpins’) are retained if log2 intensity at the T0 timepoint was > 9. Fold-changes were calculated independently for each replicate at each timepoint, and a Bayes Factor was calculated using BAGEL [32] on all replicates at each timepoint. Bayes Factors are summed across timepoints for a final gene-level Bayes Factor for each cell line.

In parallel, RNA-seq on each cell line was processed using Tophat v2 [44] and Cufflinks v2 [45] in quantitation-only mode, and later filtered for protein coding genes. Gene expression values for all cell lines were combined into a matrix of log2(FPKM + 0.01) values and the matrix was quantile normalized. The modal expression value was then calculated as the peak of the main distribution of biologically relevant gene expression (log2 FPKM = 3.76). Quantile normalized logFPKM values are in Supplementary Table 2.

### Calculating functional enrichment of correlated essentiality profiles

For all gene pairs, a Pearson correlation coefficient was calculated comparing the vectors of Bayes Factors across the 112 cell lines. Correlations were rank ordered and binned into groups of 1,000. For each bin, the enrichment for co-functionality was evaluated as the fraction of gene pairs involved in the same biological process (‘true positives’) relative to the fraction of gene pairs involved in different biological processes (‘false positives’). Biological process annotations were taken from KEGG pathways downloaded from the Molecular Signatures Database (file c2.curated.kegg.v4.0.symbols.gmt, [46]), after removing the ribosome, proteasome, and all terms with > 200 genes.

To maximize functional enrichment, three normalization and filtering methods were applied. First, each Bayes Factor for each gene in each cell line was divided by the number of hairpins used to calculate the BF. Second, the matrix of hairpin-normalized BFs was quantile normalized. This hairpin- and quantile-normalized BF is referred to as the “essentiality score”. Third, a gene was only retained for downstream analysis if it had an essentiality score > 2 (corresponding to ∼15% FDR) in at least two cell lines where the gene was also expressed at above modal expression. The filtered gene set includes 2,883 genes.

To create the coessentiality network, the mcl clustering algorithm was applied to sets of highly correlated genes using a sampling strategy across three parameters. First, correlation thresholds of 0.1%, 0.2%, 0.5%, and 1.0% FDR were applied (n=1,568, n=2,240, n=3,684, n=5,690 gene pairs respectively). Second, native correlations or correlations raised to the 4^th^ power were considered. Third, the –I parameter of mcl was applied in a range from 1.8 to 4.1 in 0.1 increments.

The output from mcl is a list of hard clusters (each gene is assigned to exactly one cluster). The functional enrichment of each mcl run output was evaluated using the same process as functional enrichment of the essentiality score correlations, but considering whether co-clustered genes were also enriched for co-functionality. The data set using correlations at FDR 0.1%, raised to the 4^th^ power, and with mcl –I 2.0 was judged to have the best combination of coverage and functional enrichment; this subset of correlations comprises the Cancer Coessentiality Network v1.0 (Supplementary Table 5) and the mcl output defines the clusters described in this study (Supplementary Table 6).

### Tumor genomic analysis

Gene-level copy number data for 481 ovarian tumors from [30] was downloaded from cBioPortal [31]. Mean log_2_ copy number across all genes in the 5q14-22 locus was calculated and samples were divided into “copy loss” (mean copy number < −0.5) and “no copy loss” (mean copy number > −0.5). Kaplan-Meier survival analysis was performed using the lifelines package in Python v 2.7.

### Validation experiments

#### Cell lines for OxPhos vulnerabilities

Cell lines were either developed in-house or kindly provided by Dr. Richard Marcotte (B. Neel Lab) or Dr. Fabrice Sircoulomb (R. Rottapel Lab) at Princess Margaret Cancer Center, and were verified by short tandem repeat (STR) profiling. A full description of each cell line used, including the tissue origin and their culture environment is detailed in Supplementary Table 9 Cell lines were selected based on their tissue subtypes, distribution in the cluster heatmap (Figure 3a), as well as the growth conditions. The representative cell lines of cluster that predicted high sensitivity to OXPHOS perturbation include five ovarian (OV1369, OV90neo, TOV1369TR, TOV1946 and TOV21G) and three breast (BT20, HCC1419 and SKBR5) cancer cell lines. The representative cell lines that are predicted to be less sensitive to OXPHOS perturbation include ten ovarian (ES2, JHOC5, OVCAR5, OVCAR8, PEA1, PEA2, PEO4, PEO6, PEO14 and SKOV3), one breast (MCF7) and two pancreatic (GP3A and MIAPACA2) cancer cell lines. To normalize for media-specific effects on cell proliferation, all cell lines were cultured in the same growth media as in the pooled shRNA screens from which the Cancer Coessentiality Network was derived (see [21]).

#### Rotenone sensitivity assays

Each cell line was plated into 15-cm culture plates (Corning, 430599) and was grown to 80% confluence in the requisite medium. Cells were washed with warm Dulbecco's phosphate-buffered saline (DPBS) (Gibco, 14190-144), treated with 0.25% trypsin-EDTA solution (Gibco, 25200-056) for five to ten minutes or until they lift off at 37°C, re-suspended in warm medium and counted using the Beckman Z2 Coulter Counter with a size gate setting between 10.00µm and 27.85µm. Cells were plated in ten 6-well plates (Corning, 3516) at ∼200,000 cells per well, in a total volume of 3mL per well. After 24 hours, the medium was replaced with 3mL of fresh medium containing 1µM rotenone or 0.1% DMSO such that each 6-well plate contains triplicates of rotenone-treated and DMSO-treated cells. A 100mM stock solution was prepared from dissolving 39.441mg of rotenone (Sigma-Aldrich, R8875) in 1mL of dimethyl sulfoxide (DMSO) Hybri-Max™ (Sigma-Aldrich, D2650). From the 100mM stock, 1mM aliquots were prepared and were stored at −20°C. Working concentration of 1µM was freshly prepared on the day of drug treatment from 1:1,000 dilution of 1mM solution in cell culture media (i.e. Final DMSO concentration of 0.1%). Cell proliferation was assessed by counting replicate populations every 12 hours for five days. Each replicate was washed with warm DPBS, trypsinized and resuspended in 2mL of DPBS. After mixing the cell suspension by gently pipetting up and down several times, 500µL of each replicate was added to 9.5mL of IsoFlow™ Sheath Fluid (Beckman Coulter, 8547008), and the prepared samples were counted using the Beckman Z2 Coulter Counter.

#### Phenoformin sensitivity assays

Cells were plated in 12-well plates (Corning, 3513) at ∼80,000-120,000 cells per well, in a total volume of 2mL per well. After 24 hours, the medium was replaced with 2mL of fresh medium containing 40µM, 100µM or 200µM phenformin or 0.1% DMSO, such that each 12-well plate contains three different concentrations of phenformin- and DMSO-treated triplicates. The working concentrations were freshly prepared from 1:1,000 dilution of 40mM, 100mM and 200mM stock solutions in cell culture media to get a final DMSO concentration of 0.1%. The cells were counted after 96 hours of proliferation in the presence of phenformin. Cells were washed with warm DPBS, trypsinized and resuspended in 1mL of DPBS. After gentle mixing of the cell suspension by pipetting up and down several times, 500µL of each sample was added to 9.5mL of IsoFlow® Sheath Fluid (Beckman Coulter, 8547008), and the prepared mixture was counted using the Beckman Z2 Coulter Counter.

#### Generation of EXO1 knockouts in ovarian cancer cell lines

Six ovarian cancer cell lines were selected based on their varying p53 status: JHOC5, OVCAR8 and TOV21G express wild-type p53; whereas OV1369, PEO6 and SKOV3 harbor three different p53 mutants including G244C, G244D and null, respectively. A full description of each cell line used, including the tissue origins of cell lines and their culture environment is summarized in Supplemental Table 10. Three sgRNAs targeting EXO1 and one sgRNA targeting LacZ were chosen from the Toronto Knockout (TKO) library [24]. The forward and reverse oligonucleotides including the 20 base pair target sequence and the BsmBI overhangs were synthesized as follows.

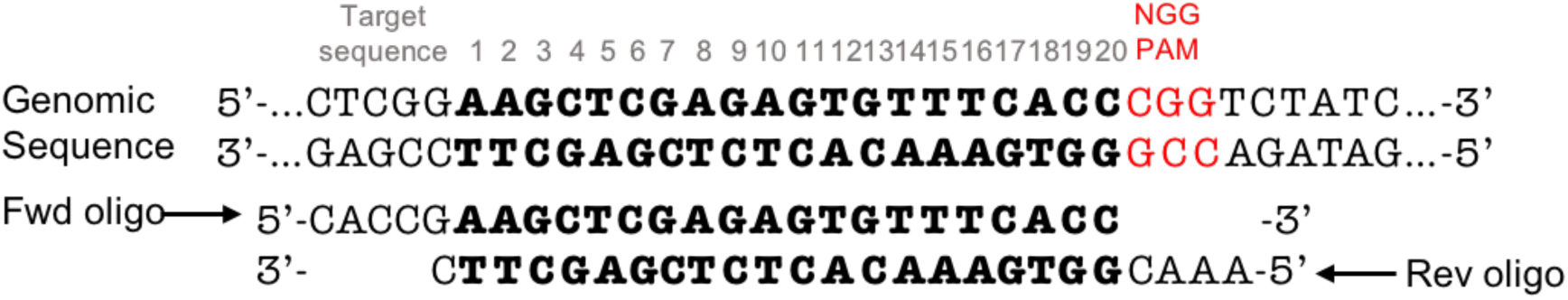

Each pair of oligos was phosphorylated and annealed. LentiCRISPRv2 (LCv2) (Addgene, plasmid #52961) [47], a single vector containing two expression cassettes, *Sp*Cas9 and the guide RNA, were digested using BsmBI, and the annealed oligos were cloned into the sgRNA scaffold. The resulting ligation products, LCv2-EXO1 and LCv2-LacZ, were transformed into One Shot® Stbl3™ chemically competent *E. Coli* (Invitrogen, C7373-03), and the plasmid DNA was isolated using the PureLink® HiPure Plasmid Midiprep Kit (Invitrogen, K2100-04).

For lentivirus production, low passage number 293T packaging cells were seeded in low-antibiotic growth media at ∼350,000 cells per 10cm plate (Corning, 430167). After 24 hours of incubation, a mixture of the three transfection plasmids, 5400ng of packaging vector psPAX2, 600ng of envelope vector pMD.2G and 6.0µg of LCv2-EXO1 or LCv2-LacZ, as well as a master mix of OPTI-MEM (540µL per transfection; Gibco, 31985-062) and X-treme GENE 9 DNA transfection reagent (36µL per transfection; Roche, 06-365-809-001) were prepared. The transfection mix was incubated for thirty minutes before added to the packaging cells. At 24 hours post-transfection, the medium was changed to remove the transfection reagent and replaced with 10mL of high-bovine serum albumin (BSA) growth medium per 10cm plate for viral harvests. The lentivirus-containing medium was harvested at ∼48 hours post-transfection and new high-BSA growth medium were added to cells for second viral harvests at ∼72 hours post-transfection. The harvested lentivirus was aliquoted in sterile 15-mL conical polypropylene centrifuge tubes (Corning, 430791) and stored at −80°C. For infection, cells were seeded in 6-well plates (Corning, 3516) at 300,000 cells per well and infected with the lentiviral sgRNA targeted to EXO1 or LacZ along with 8µg/mL polybrene (Sigma, H9268) to increase the virus infection efficiency. After 24 hours, the virus-containing medium was removed and replaced with new medium containing 2µg/mL puromycin for selection of transduced cells. Cells were incubated for an additional 48 hours or until the no-infection control is completely wiped out. Each of the puromycin-selected infected populations was expanded to 10-cm culture plates.

Stable EXO1 knockout (EXO1-KO) lines were generated by transducing a lentiviral construct (LCv2) containing Cas9 nuclease and sgRNA targeting EXO1 into six ovarian cancer cell lines of varying p53 status: JHOC5 (wild-type), OVCAR8(wild-type), TOV21G (wild-type), OV1369 (G244C), PEO6 (G244D) and SKOV3 (null). Following the lentiviral infection, puromycin was used for selection of stably transduced cells, and knockouts were confirmed by western blot.

#### Cell proliferation assays with Nutlin-3a

Cells were seeded in 24-well plates at ∼50,000 cells in a volume of 1mL per well, or in 12-well plates at ∼100,000 cells in a volume of 2mL per well. After 24 hours, the medium was replaced with 1 or 2mL of fresh medium containing 10µM nutlin-3a or 0.1% DMSO in triplicates. Working concentration of 10µM was freshly prepared on the day of drug treatment from 1:1,000 dilution of 10mM stock solution in cell culture media (i.e. Final DMSO concentration of 0.1%). Cell proliferation was assessed by counting replicate populations after 96 hours of drug treatment. Each replicate was washed with warm DPBS, trypsinized and resuspended in 1 or 2mL of DPBS. After mixing the cell suspension by gently pipetting up and down several times, 500µL of each replicate was sampled for counting with the Beckman Z2 Coulter Counter.

